# Itinerant complexity in networks of intrinsically bursting neurons

**DOI:** 10.1101/2020.03.22.002170

**Authors:** Siva Venkadesh, Ernest Barreto, Giorgio A. Ascoli

## Abstract

Active neurons can be broadly classified by their intrinsic oscillation patterns into two classes characterized by periodic spiking or periodic bursting. Here we show that networks of identical bursting neurons with inhibitory pulsatory coupling exhibit itinerant dynamics. Using the relative phases of bursts between neurons, we numerically demonstrate that the network exhibits endogenous transitions among multiple modes of transient synchrony. This is true even for bursts consisting of two spikes. In contrast, our simulations reveal that identical singlet-spiking neurons do not exhibit such complexity in the network. These results suggest a role for bursting dynamics in realizing itinerant complexity in neural circuits.

## I. Introduction

Neural systems exhibit transitions across multiple spatiotemporal scales. While individual neurons exhibit spiking events, which are sharp changes to the resting membrane potential, an ensemble of self-organized neurons can exhibit transitions among multiple coexisting metastable states [1–3]. In a metastable state, the interacting elements enter into a transiently fixed relationship to each other, or “mode”, before subsequently diverging and visiting a different transient mode. Metastability has been suggested to underlie the necessary coordination within and between brain regions [2]. Moreover, experimental evidence shows that the coordinated transitions among neural ensembles correlate with changes in an organism’s behavior [4–8].

Chaotic itinerancy [9] is a special case of metastability and has been observed in many complex systems, including globally coupled maps [10,11] and electrically coupled point neurons [12–14]. Those coupled systems show endogenous transitions through a sequence of quasi attractors in the state space. Among possible scenarios underlying itinerancy in coupled systems [15] are attractors with riddled basins [16], where initial conditions that are arbitrarily close to an attractor can generate trajectories leading to a different attractor. However, the intrinsic property of interacting elements that is necessary for the emergence of network itinerancy has not yet been clarified.

Neurons can be dynamically classified based on their intrinsic patterns of activation [17]. In the current work, we constructed three separate networks of identical elements based on two broad classes of neurons: bursting neurons, which have either chaotic or two-loop periodic trajectories in the phase space, and spiking neurons, which have a single-loop trajectory. Analysis of the ensuing network dynamics revealed several novel results. First, networks of bursting neurons showed transiently stable phase differences in pairs of neurons at the level of bursts, which transitioned to other transiently stable arrangements endogenously. Previous reports have described ensembles of bursting oscillators that synchronize on the bursting timescale and desynchronize on the spiking timescale [18]. Our results show that synchronized bursting is only one of the *multiple* modes of stability observed in the network. We illustrate these complex dynamics using the burst-level phase differences between neurons as a variable of coordination. Second, we demonstrate that even networks consisting of bursting elements as simple as doublet-spikers can exhibit multiple stable phase differences and endogenous transitions. Third, singlet-spiking neurons only display, in contrast, perfectly phase-synchronized and desynchronized modes between pairs of neurons in the network. These results suggest that complex periodic oscillators such as bursters are crucial for the emergence of itinerant dynamics in neural networks.

## II. Burst-level phase difference as a coordination variable

We use Izhikevich model (IM) neurons [19], whose dynamics are described by (eqn1-2):

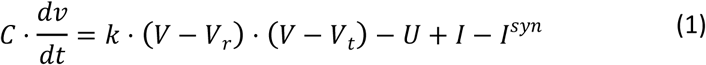

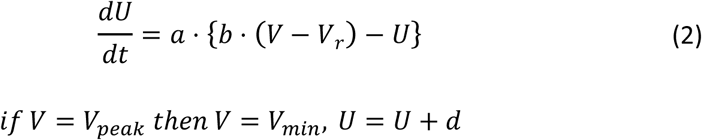

Here, *I* is a constant current, and *I*^*syn*^ is the total synaptic current from all presynaptic neurons. The parameters were identified by an optimization framework [20] to match the spike patterns of an isolated (*I*^*syn*^=0) IM neuron to experimentally-obtained voltage traces from a stuttering CA1 neurogliaform cell (a GABAergic neuron type) from the rodent hippocampus [21] (*k* = 3.59, *a* = 0.01, *b* = −10, *d* = 120, *C* = 195, *V*_*r*_ = −63.5, *V*_*t*_ = −46.6, *V*_*peak*_ = 11.4, *V*_*min*_ = −50.6). The bifurcation diagram with respect to the constant input current *I* was obtained by making a Poincaré cut at *V* = *V*_*peak*_ − 20*mV* in the direction of increasing *U* after discarding 1s of initial transient behavior. This revealed period-doubling cascades leading to chaos (Fig 1A). For chaotic bursting neurons, we set *I*=500*pA.*

**Fig 1.**
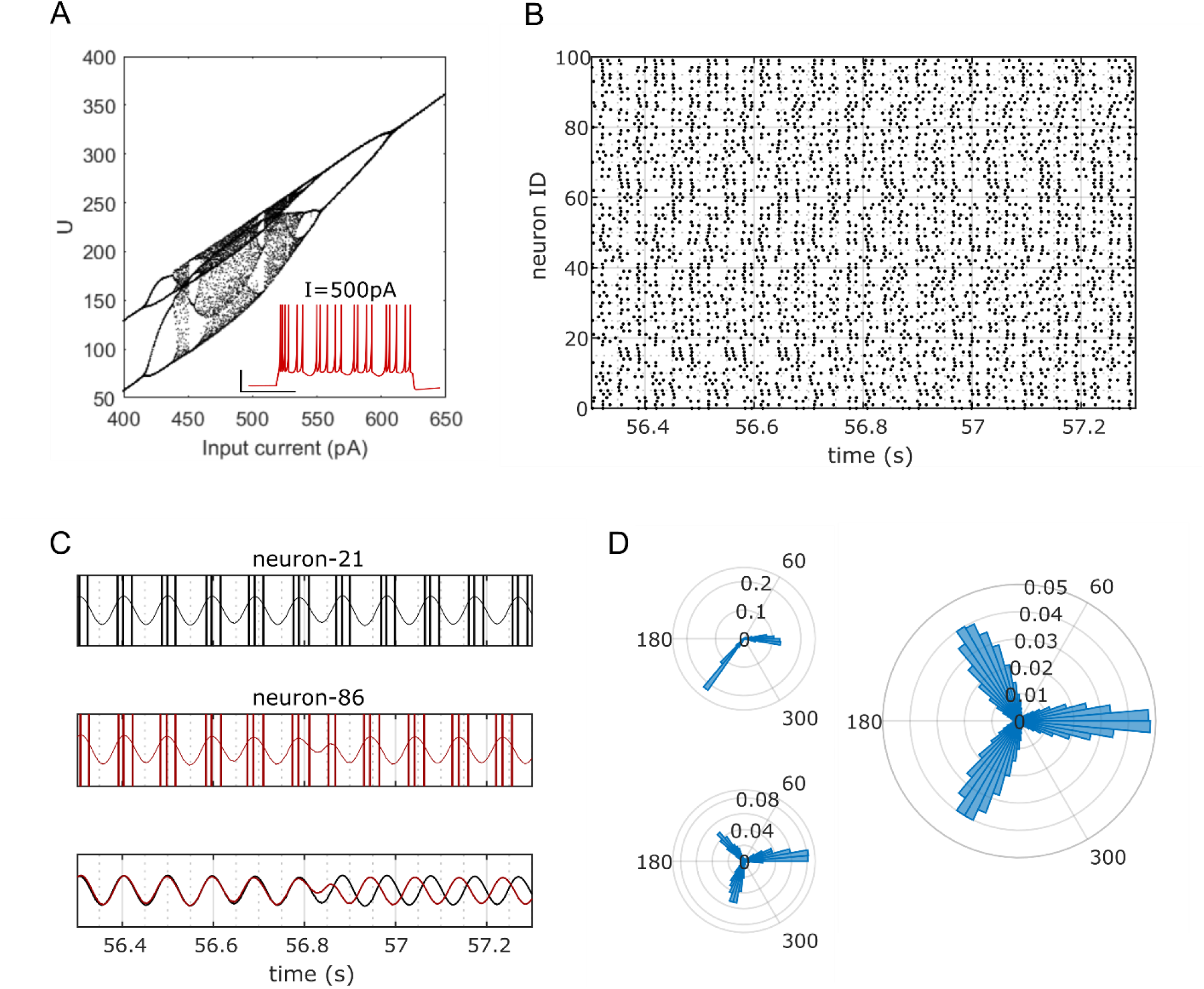
Endogenously transitioning phase-locked modes in a network of 100 identical bursting neurons. **A.** Bifurcation diagram of the bursting neuron showing period-doubling cascades leading to chaos for increasing input current. Inset shows the activity pattern (voltage vs. time) of the isolated neuron. **B.** Raster plot showing the spike times of a network of neurons for a duration of 1 second. **C.** The discrete spike times of two neurons are transformed into a continuous signal corresponding to burst cycles (top & middle, also see Fig S1). The phase-locked mode endogenously transitions from mode − 0 to mode − 4*π*/3 (bottom). **D.** Normalized distributions of phase differences between the neurons shown in C during the same 1 second window as in C (left-top), and for 115 seconds (left-bottom). The distribution of phase differences between 100 randomly selected neuron pairs (right) show three clearly preferred locked modes (0, 2*π*/3, 4*π*/3).

A network of 100 identical neurons, which were coupled using a delta function (eqn-3), was then constructed with

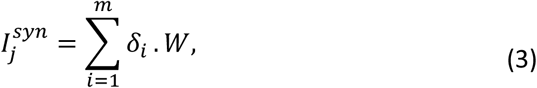

where *m* is the number of presynaptic neurons connecting to the postsynaptic neuron *j* and *W* is the inhibitory connection weight. *δ*_*i*_ is 1 (for a single time step) if neuron *i* spikes and is 0 otherwise. All networks were constructed with a connection probability of 0.7 and a constant *W* (*W* = 8 unless specified otherwise). Simulations were performed with CARLsim [22] for a duration of 120s using the fourth order Runge-Kutta integration method with 100 steps per millisecond. The first 5s of simulation was discarded, and the analysis was performed for a total duration (Δ*T*) of 115s.

In order to extract the relative phases of bursts, the following steps were carried out for each neuron. First, discrete spike events were lowpass-filtered to obtain a continuous periodic signal which captures bursting cycles (Fig 1C, also see Fig S1A-C top). Then, the instantaneous phase of this signal was extracted using the Hilbert transform. Finally, the instantaneous phase differences between pairs of neurons were calculated for each millisecond in a given duration. We find that pairs of neurons predominantly exhibit phase differences near 0, 2*π*/3, or 4*π*/3 radians, and that they endogenously transition among these transiently locked states (Fig 1D).

Next, we studied the stability of transiently phase-locked modes and the nature of transitions among them. Specifically, we asked the following questions pertaining to pairs of neurons: (i) What is the locked duration in each of the three modes? (ii) For what fraction of the total duration (Δ*T*) does a pair exhibit stable phase differences in each of the three modes? (iii) What are the transition probabilities among the three modes?

The stability of a pair of neurons over a duration *δT* was quantified by averaging all the instantaneous phase differences Δ*θ*(*t*) represented on a unit circle (eqn-4). *δT* = 500*ms* unless specified otherwise.

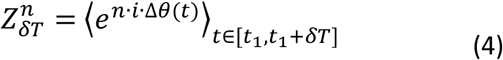

Here, ⟨*f*⟩ denotes the time average of *f*. The magnitude of 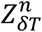 was used as a measure of stability. The parameter *n* indicates the number of stable clusters of phase differences between pairs of neurons. For example, if the bursts of two neurons are perfectly synchronized over *δT*, then 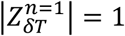. If their phase differences segregate into two equally populated clusters at 0 and *π* radians, then 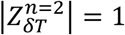 and 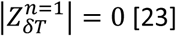 [23].

A pair of neurons was deemed to be ‘stable’ or ‘phase-locked’ in a mode if 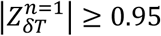 during *δT*. Such stable pairs were also assigned a mode 0, 2*π*/3, or 4*π*/3 based on the angular range in which the phase of 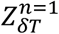 lies (see Fig 1D for the bounds of the three ranges). In order to identify how long a pair remains phase-locked in a given mode, 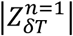 was calculated sequentially for non-overlapping *δT*s until either the pair was unstable (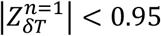 for a duration *δT*) or a new mode was detected. The sum of all stable intervals (*δT*s) calculated from this sequential search was taken as the locked duration of the given mode. This gives a sequence of modes and a locked duration (in increments of *δT*) in each mode visited by a pair.

The locked durations of any of the three modes for 100 randomly selected pairs were exponentially distributed (Fig 2A-C), consistent with a previous report [11]. Pairs exhibit an expected locked duration of roughly 2*s* in all three modes. It is worth mentioning here that the expected locked durations do not give information about the fraction of the overall time Δ*T* that such locked modes were observed, because of their transient nature. We refer to this fraction as the fraction of locked time, given by *N* × *δT*/Δ*T*, where *N* is the total number of phase-locked *δT* intervals observed for a pair. The average of the fractions of locked time was 0.22 for each mode (Fig 2A-C insets).

**Fig 2.**
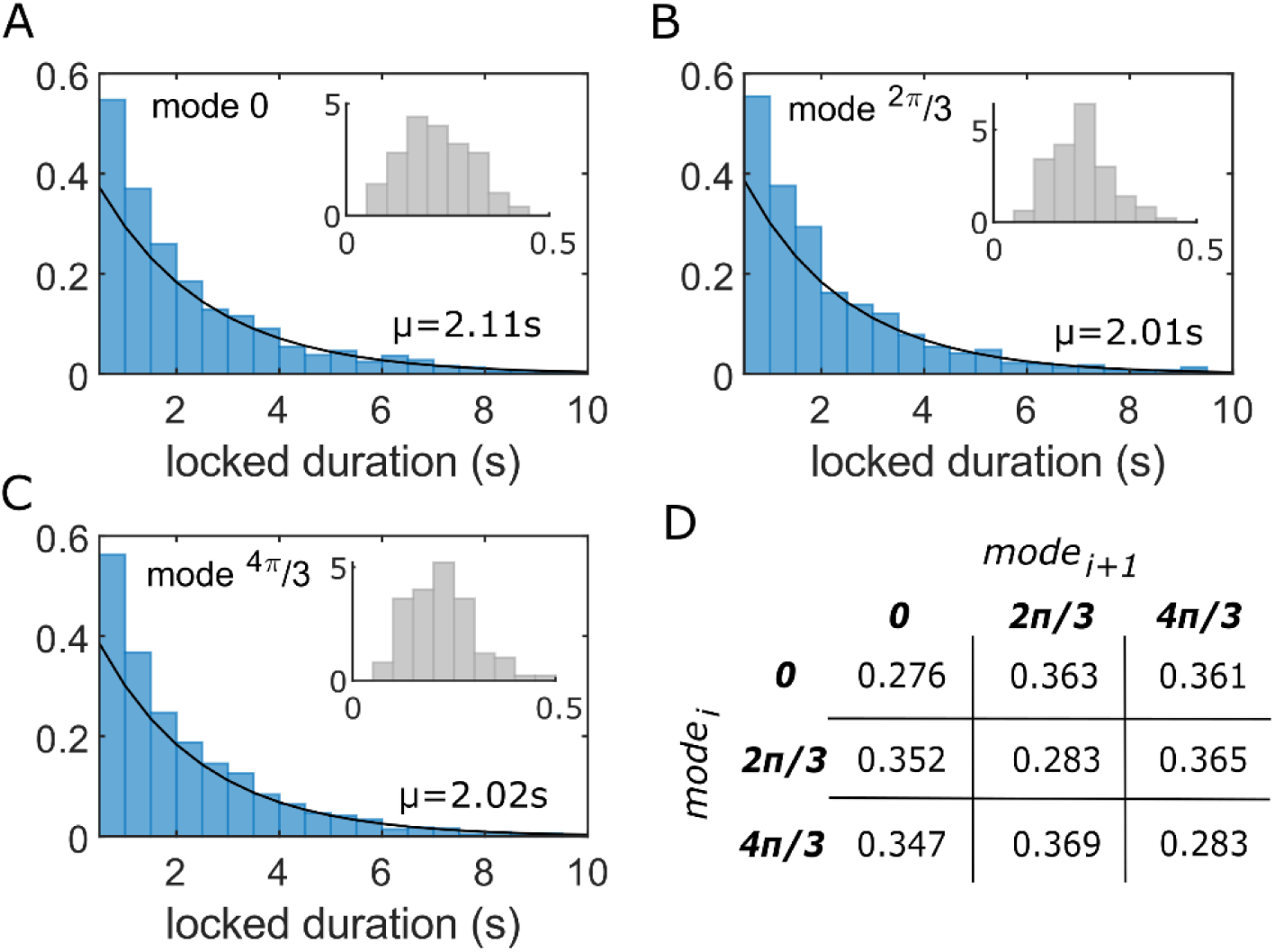
Stability and transitions in the network of chaotic bursting neurons. **A-C.** Probability densities of locked durations in modes − 0 (A), 2*π*/3 (B) and 4*π*/3 (C) are exponentially distributed with the expected locked durations (*μ*) of 2.11s, 2.01s and 2.02s with 95% confidence intervals [2.0, 2.24], [1.91, 2.13] and [1.91, 2.14] respectively for 100 randomly selected pairs. Insets show probability density distributions of the fractions of locked time for 100 pairs. **D.** Probabilities of transitions among the three modes. Each table entry denotes the probability of a pair transitioning from *mode*_*i*_ to *mode*_*i*+1_ after losing its stability from *mode*_*i*_, where i denotes the sequence index.

In addition, the sequences of modes visited by 100 pairs were used to gain insights into the nature of the transitions. The counts of transitions among the modes, which were given by the number of pairwise occurrences of modes in the sequence, were used to construct a 3×3 mode transition probability matrix (Fig 2D). Interestingly, there were several occurrences where pairs of neurons, after becoming unstable, failed to exhibit successful transitions. In other words, after losing stability from a certain locked mode, they were attracted back to the same mode rather than escaping to a different one (see Fig S2 for an example). The probability of successfully transitioning to a different mode (i.e. escape probability) is 0.72 from each of the three modes (Fig 2D). The apparently random nature of successful transitions from a given mode, which is noted by the approximately equal probabilities of transitions to the *other* two modes, is suggestive of a high entropic network exhibiting states that persist over several cycles of bursts.

## III. Itinerancy in the simplest form of bursting

Bursting dynamics include a spectrum of oscillatory patterns ranging from a double loop trajectory in the phase space (e.g. doublet-spiking observed for *I*=580pA in Fig 1A) to an aperiodic trajectory (*I*=500pA in Fig 1A). In this section, we consider networks separately consisting of doublet-spiking neurons (the simplest form of bursting with only two timescales) and singlet-spiking neurons. While the periodicity of oscillations in the chaotic-spiking IM neuron depends on the bifurcation parameter *I* (Fig 1A), here we considered IM neurons with the highest periodicity of only two for doublet-spiking and one for singlet-spiking (Fig 3A).

**Fig 3.**
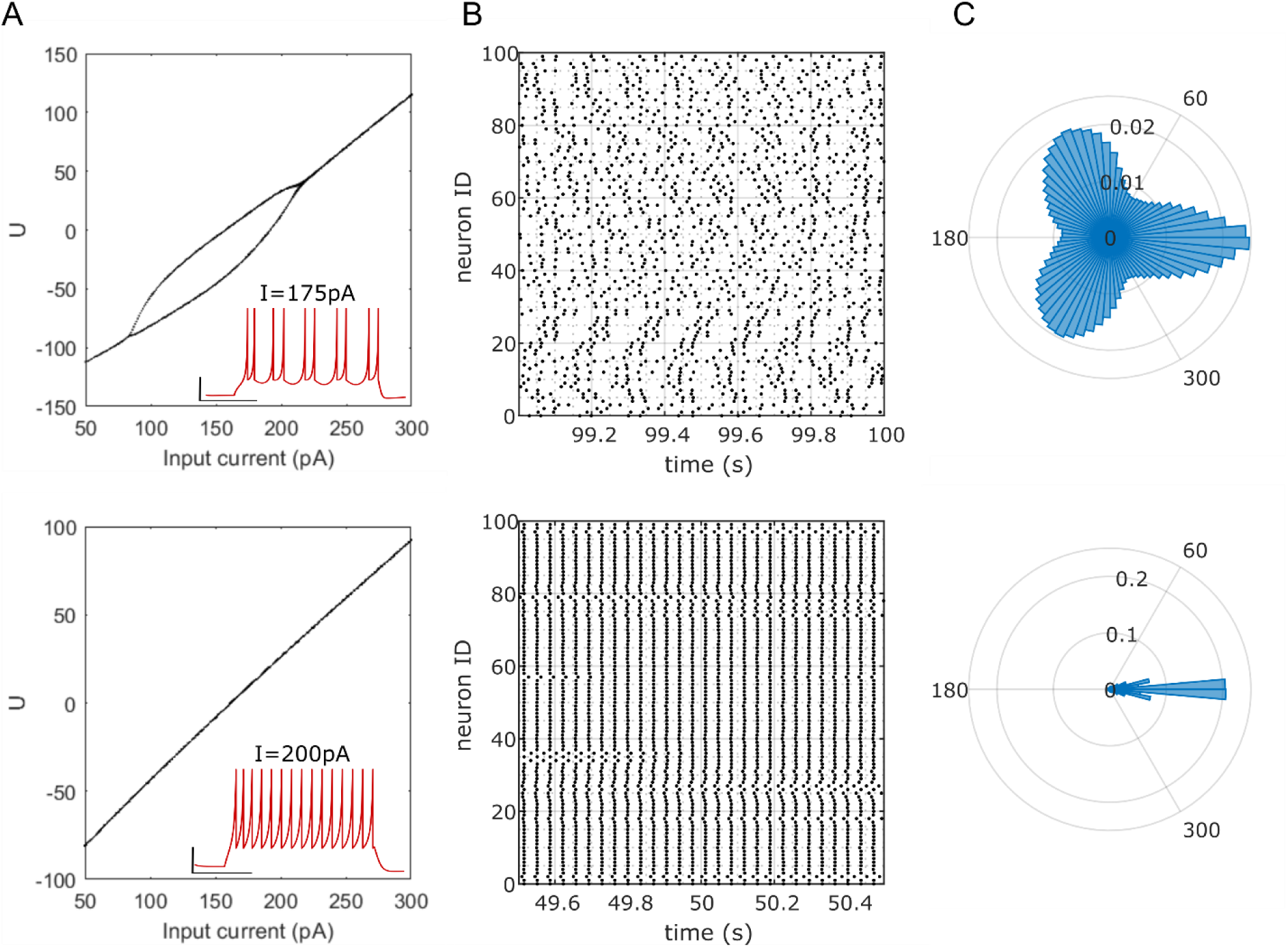
Dynamical complexity is preserved in the network of doublet- (ND) spiking neurons. **A.** An isolated neuron with a 2-loop limit cycle attractor (spike doublets shown in inset) was obtained by reducing the value of parameter *k* from the chaotic-model to 1.5 (top). Further reducing the value of *k* to 0.5 result in a singlet-spiking model (bottom). **B**. Raster plot shows the spike times of ND (top) and NS (bottom) in the network for a duration of 1 second. **C.** The phase differences between 100 randomly selected neuron pairs in the ND (top) show three preferred locked modes (0, 2*π*/3, 4*π*/3), although these distributions have a wider spread compared to NC (see Fig 1). There is only a single preferred locked mode for NS (bottom), although they show scattered distributions of non-zero phase differences (see Fig S3).

It was previously reported that bursting dynamics do not exist for low values of the parameter *k* in the IM neuron (see eqn-1) [24]. We obtained doublet- and singlet-spiking neuron models by only varying parameter *k* from the chaotic-spiking model (*k*=1.5; *I*=175*pA* and *k*=0.5; *I*=200*pA* for doublet- and singlet-spiking neurons respectively) (Fig 3). It should be noted that the bifurcation diagrams only illustrate the asymptotic behavior of the isolated neurons (Figs 1A&3A). Networks were constructed using these models and the steps explained in the previous section were carried out for the analysis (Fig S3, also see S1A-C middle & bottom). We found that the network of doublet-spikers (ND) showed phase-differences that were clustered near 0, 2*π*/3, and 4*π*/3 radians (Figs 3B-C top & S3A-B) like the network of chaotic-spikers (NC). However, the most stable network was realized at *W* = 20 for ND. Interestingly, the network of singlet-spikers (NS) only phase-locked near 0 radians (Figs 3B-C bottom & S3C-D).

The average locked duration in the ND is less than a second for each of the three modes (Fig 4A). The average fractions of locked times are 0.1, 0.07 and 0.07 (Fig 4A inset), and the escape probabilities are 0.62, 0.66, and 0.67 for the modes 0, 2*π*/3, and 4*π*/3 respectively. Thus, the network of neurons that are intrinsically as simple as doublet-spikers exhibits itinerant dynamics. Although the NS did not display multiple metastable phase differences, its neuron pairs still demonstrated simpler metastability, where desynchronized spiking occurred in between synchronized (mode 0) spiking (Fig S3 C-D). This is similar to the transitory dynamics reported in [12–14], where networks of point neurons coupled with gap junctions alternated between synchronized and desynchronized states. The NS showed an average locked duration of over 5s and an average fraction of locked time of 0.74 in mode 0 (Fig 4C left). While the transitions among the three metastable modes in NC and ND are generally abrupt (for instance, see Fig 1C), the transitions between successive synchronized states in NS showed scattered phase differences (Fig S3C). The occasionally slower dynamics of such transitions lead to a few stable *δT*s in mode 2*π*/3. However, the fraction of locked time in mode 2*π*/3 is negligible (Fig 4C right, also see Fig 3C bottom).

**Fig 4.**
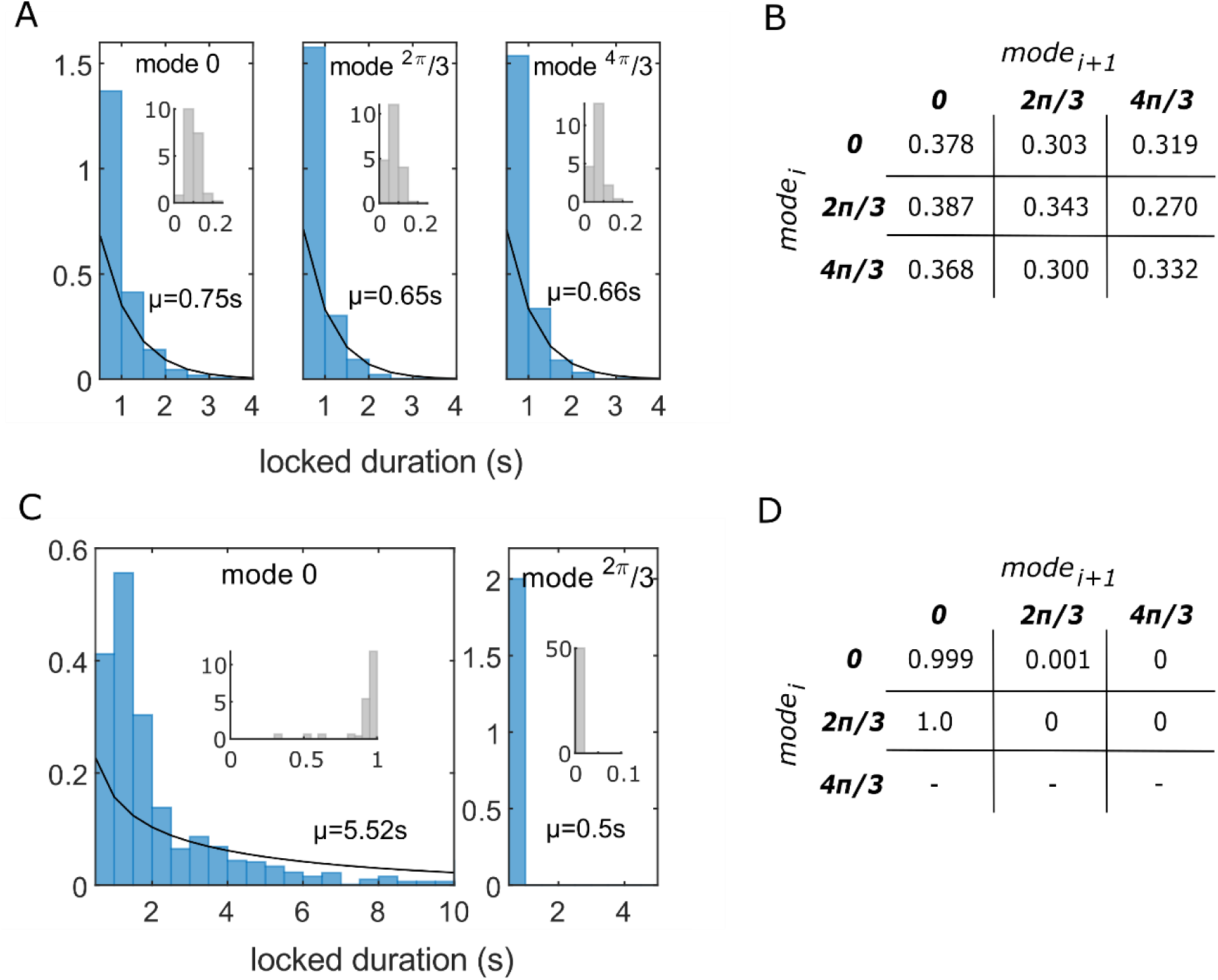
Stability and transitions in network of doublet- (ND) and singlet- (NS) spikers. **A.** Distributions of locked durations in modes − 0 2*π*/3 and 4*π*/3 are exponentially distributed in ND with the expected locked durations (*μ*) of 0.75s, 0.65s and 0.66s with 95% confidence intervals (CI) in [0.71, 0.79], [0.61, 0.68] and [0.63, 0.70], respectively, for 100 randomly selected pairs. **B.** Probabilities of transitions among the three modes in ND. **C.** The locked durations in mode − 0 (left) roughly follow a gamma distribution in NS with *μ* = 5.52*s* (*α* = 0.54 with CI [0.51, 0.57] and *β* = 10.29 with CI [9.41, 11.25] are the shape and scale parameters, respectively, of the gamma distribution, and *μ* = *α* × *β*). There are 2 occurrences of locked mode − 2*π*/3 (right), which are due to the slower phase scattering dynamics. Insets show probability density distributions of the fraction of total time a pair was locked in the mode. **D.** Probabilities of transitions in NS show no stable transitions to the mode − 4*π*/3.

The itinerant dynamics that emerges from the collective interaction of complex-periodic spiking neurons is also sensitive to the network connectivity parameters such as *W*. Therefore, we studied how the tri-stability (as illustrated in Fig 1D – right and Fig 3C – top) is affected by changing the strength of the inhibitory connections. We plotted two measures, 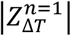 and 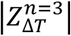 (see eqn – 4) for connection weights in [0, 40]. For a weakly connected NC (*W* = 4), both measures are near zero (Fig 5A – left) because the phase differences between pairs of neurons are mostly asynchronous and they are uniformly distributed with predominantly transitioning *δT*s (Fig 5B – top). At *W* = 8, 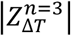 reaches its maximum value of 0.62, while 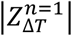 remains near zero. These measures correspond to the tri-stability illustrated in Fig 5B – middle, where the stable *δT*s are more prevalent than the transitioning ones. Similarly, the maximum of 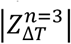, when the corresponding 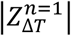 is near zero, is only 0.22 for the ND (Fig 5A – middle), as their phase differences are more spread-out (Fig 5C – bottom) than the most stable NC. Furthermore, the ND required stronger inhibitory connections (*W* = 20) to reach this maximum (Fig 5A – middle). Interestingly, for weaker connections (*W* = 4), ND’s metastability is qualitatively similar to that of the NS, where 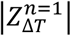 scores higher than 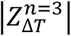 (Fig 5A – middle & right), and the modes 2*π*/3 and 4*π*/3 are nearly non-existent (Fig 5C – top & Fig 5D).

**Fig 5.**
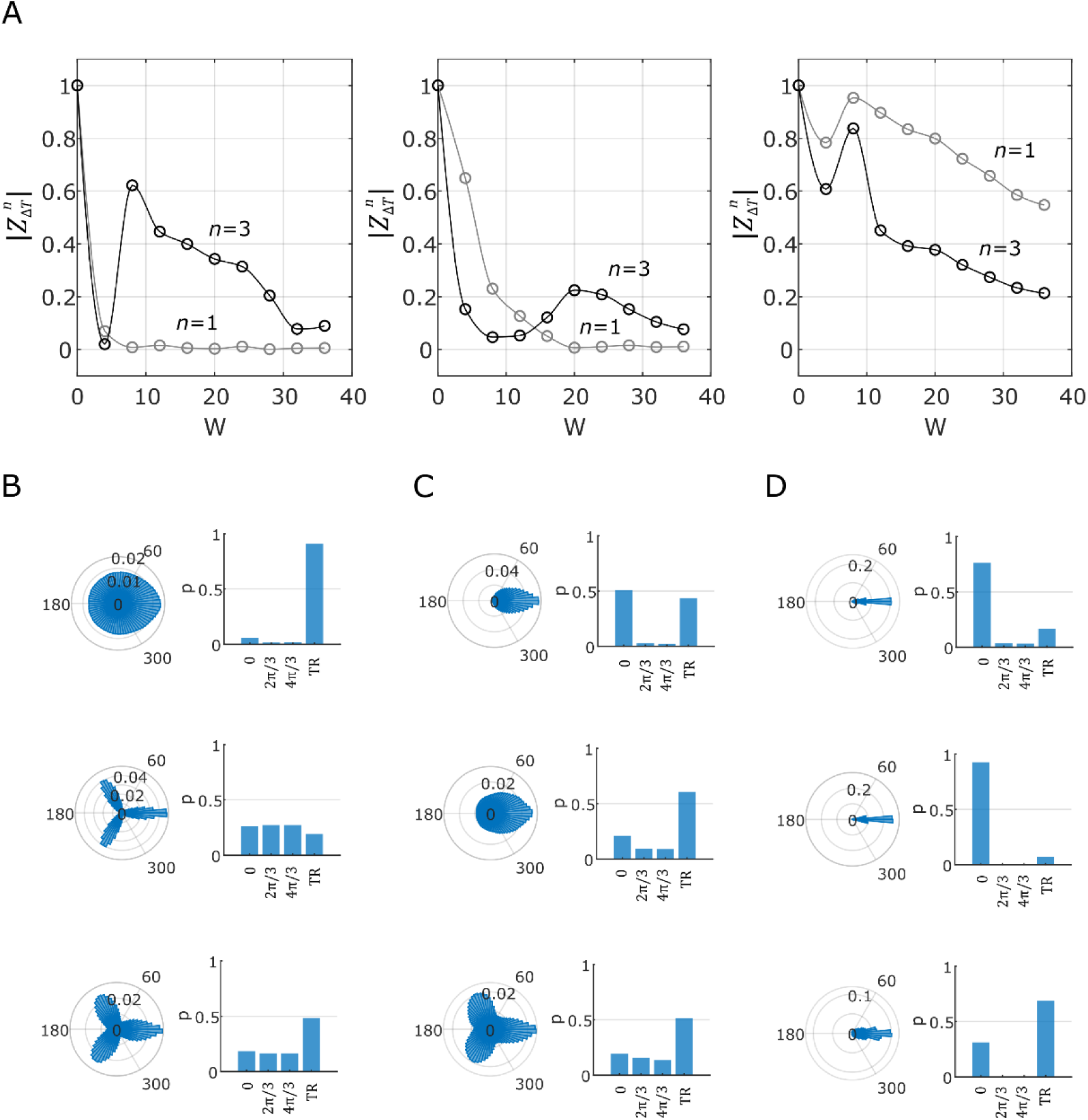
Stability as a function of connection weight. **A.** Magnitude of the average of phase differences between 100 randomly selected pairs of neurons (see eqn – 4) for NC (left), ND (middle) and NS (right). Δ*T* = 115*s*. Note that *W* = 0 corresponds to isolated neurons that fire in perfect synchrony. Therefore, 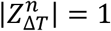 for any *n*. **B.** Distributions of phase differences for weakly connected NC (top-left) and distributions of the probabilities of the modes visited (top-right) Here, *W* = 4. A total of 1000 *δT*s was analyzed to calculate the probabilities. *δT* = 100*ms* and ‘TR’ denotes transitioning *δT*s. **B – D:** Distributions of phase differences and the probabilities of visited modes when *W* = 4 (top), *W* = 8 (middle) and *W* = 20 (bottom) for the NC (B), ND (C) and NS (D).

Thus, there are optimal connection weights that maximize the stability of the states visited by the networks of bursters. In addition, the network of simpler bursters exhibits states that are less stable than those of the network of more complex bursters.

## IV. Summary and discussion

Metastability is a useful framework to characterize the existence of synchronized and desynchronized states in a system [2,25]. However, the relationship between metastability and itinerancy has not been very clear. It has been suggested that the appearance of metastable states is a necessary but not sufficient condition for itinerancy [26]. Along this line, a few scenarios for the existence of itinerancy have been proposed [16]. The current study finds that while the networks of both the simple-periodic (singlet-spiking) neurons and complex-periodic (chaotic- and doublet-spiking) neurons show metastability, only the latter class showed *multiple* synchronized modes.

The chaotic-spiking model used in this work was obtained by tuning the IM parameters to reproduce the stuttering behavior of a neurogliaform interneuron [21]. Neurogliaform interneurons in the cerebral cortex connect nonspecifically to almost all other neuron types within their somatic layer as well as across layers [27]. In addition to electrical and chemical synapses, they also influence target neurons by volume release of GABA and were suggested to play a crucial role in broadly regulating the synchronized activity of neural circuits [21,28,29], a role aptly described as “master regulators” [27].

The network of identical simple-periodic spiking neurons did not exhibit itinerant complexity. However, it should be noted that the broad class of such simple-periodic neurons include a range of spiking timescales such as the regular spiking observed in many pyramidal neurons and fast-spiking observed in many Parvalbumin positive interneurons near their respective rheobases. A system can exhibit bursting if it consists of mutually interacting fast and slow subsystems, where the slow dynamics modulate the fast-spiking [30]. Further work is required to determine if the additional mechanisms that are necessary to induce bursting in simple-periodic neurons are sufficient for the emergence of itinerant complexity.

## Code availability

Network simulation and analysis scripts are available at https://github.com/sivaven/Metastability.git

## Acknowledgments

We thank Dr. Bard Ermentrout for his feedback in the early stages of this project. We also thank Dr. Ken De Jong and Dr. Jeff Krichmar for their feedback in the final stages of this project. This work was supported in part by National Institutes of Health (NIH) grants R01NS39600 and U01MH114829.

