## Supplementary figures for "Itinerant complexity in networks of intrinsically bursting neurons"

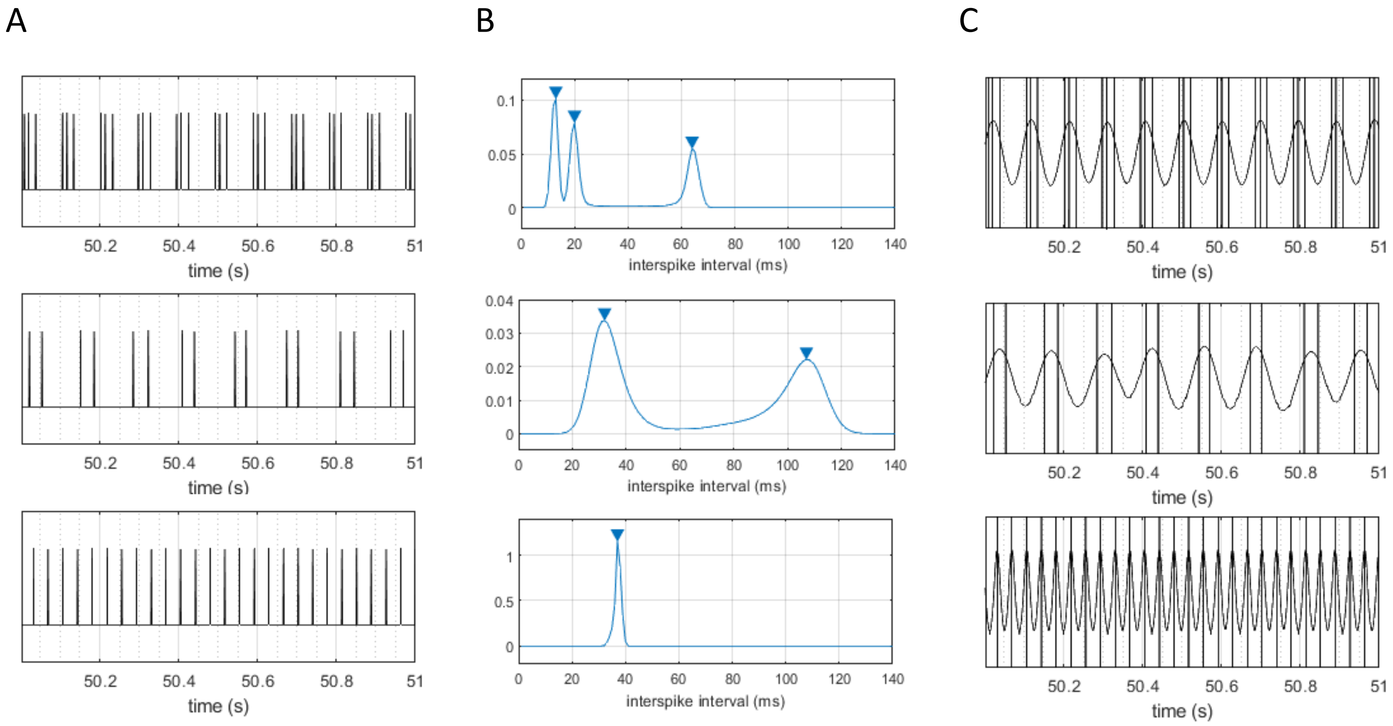


**Fig S1**. Transforming discrete spike times into a continuous signal capturing bursting cycle. **A.** Activity patterns of a sample neuron for a duration of one second, which show the existence of periodic bursting in NC (top) and ND (middle) and periodic spiking in NS (bottom). **B.** Probability density estimates of interspike intervals from all the neurons. The sum of all the local maxima (triangles) are 97, 139 and 37 respectively for NC (top), ND (middle), and NS (bottom). **C.** Lowpass filtering of spikes to extract bursting cycles using Gaussian convolution. The length of the Gaussian window is equal to the duration of one full cycle of a burst (NC and ND) or a spike (NS), which was estimated from B. This is done so that any frequency higher than that of the burst frequency is filtered in NC and ND.

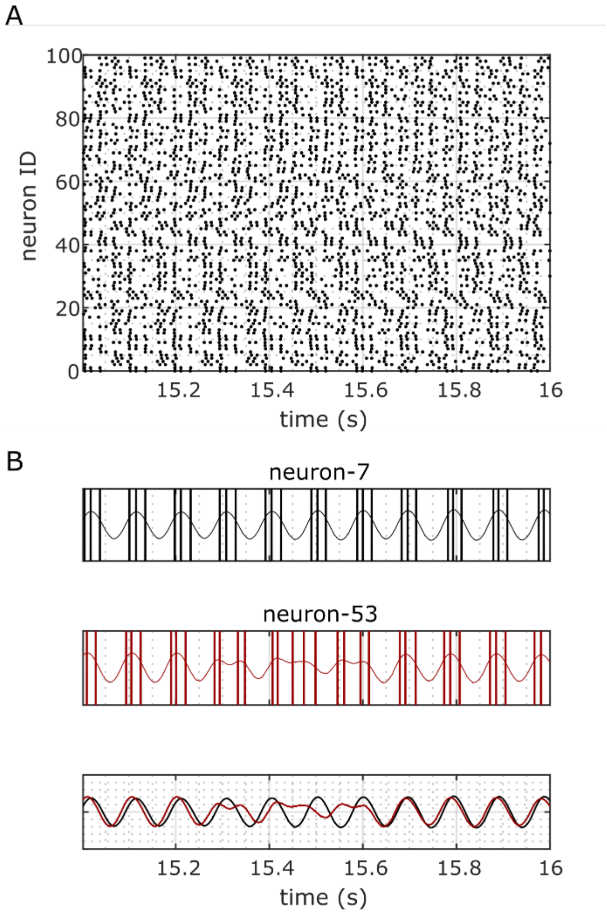


**Fig S2.** An example of an unsuccessful transition. Neuron-53 was attracted back to a relative phase near 0 radians after losing stability from the same with respect to neuron-7.

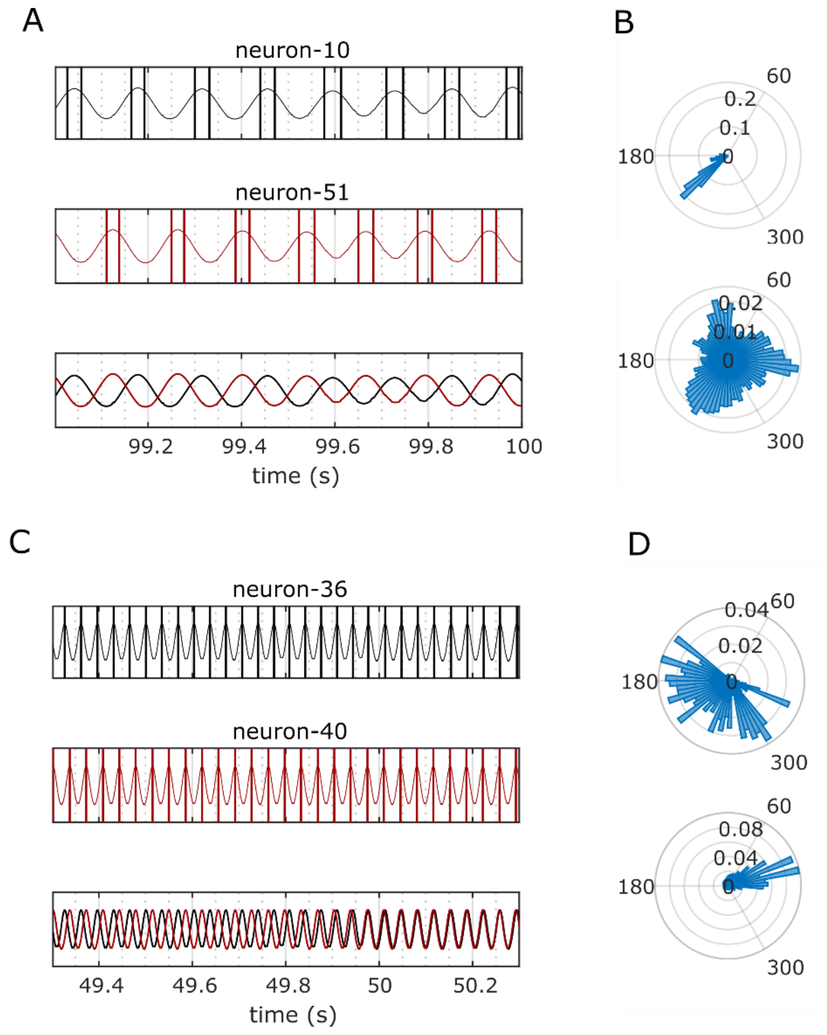


**Fig S3.** Phase-locked modes in ND and NS. **A&C.** Discrete spike times of two neurons are transformed into continuous signals corresponding to burst cycles for ND (A) and spike cycles for NS (C). **B.** The neuron pair in A shows a locked mode at $4\pi/3$ radians for 1 second (top) and its distribution for 115 second duration is shown in bottom. **D.** The neuron pair in C shows scattered phase differences for the first 500ms (top), but overall it shows a preferred locked mode near zero radian for 115 second duration (bottom).
